# Molecular changes during extended neoadjuvant letrozole treatment of breast cancer: distinguishing acquired resistance from dormant tumours

**DOI:** 10.1101/426155

**Authors:** Cigdem Selli, Arran K. Turnbull, Dominic A. Pearce, Ang Li, Anu Fernando, Jimi Wills, Lorna Renshaw, Jeremy S. Thomas, J. Michael Dixon, Andrew H. Sims

**Author notes:** Corresponding author: Andrew H Sims, PhD.

## Abstract

**Background:** The risk of recurrence for endocrine-treated breast cancer patients persists for many years or even decades following surgery and apparently successful adjuvant therapy. This period of dormancy and acquired resistance is inherently difficult to investigate, previous efforts have been limited to *in vitro* or *in vivo* approaches. In this study, sequential tumour samples from patients receiving extended neoadjuvant endocrine treatment were characterised as a novel clinical model.

**Methods:** Consecutive tumour samples from 62 patients undergoing extended (4-45 months) neoadjuvant aromatase inhibitor, letrozole, therapy were subjected to transcriptomic and proteomic analysis, representing before (≤0), early-on (13-120 days) and long-term (>120 days) neoadjuvant letrozole treatment. Patients with at least a 40% initial reduction in tumour size by 4 months of treatment were included. Of these, 42 patients with no subsequent progression were classified as “dormant”, and the remaining 20 patients as “acquired resistant”.

**Results:** Changes in gene expression in dormant tumours begin early and become more pronounced at later timepoints. Therapy-induced changes in resistant tumours were common features of treatment, rather than being specific to resistant phenotype. Comparative analysis of long-term treated dormant and resistant tumours highlighted changes in epigenetics pathways including DNA methylation and histone acetylation. DNA methylation marks 5-methylcytosine and 5-hydroxymethylcytosine were significantly reduced in resistant tumours compared to dormant tissues after extended letrozole treatment.

**Conclusions:** This is the first patient-matched gene expression study investigating long-term aromatase inhibitor-induced dormancy and acquired resistance in breast cancer. Dormant tumours continue to change during treatment whereas acquired resistant tumours more closely resemble their diagnostic samples. Global loss of DNA methylation was observed in resistant tumours under extended treatment. Epigenetic alterations may lead to escape from dormancy and drive acquired resistance in a subset of patients supporting a potential role for therapy targeted at these epigenetic alterations in the management of endocrine resistant breast cancer.

## Background

Approximately 70% of breast cancer patients who have oestrogen receptor alpha (ER) positive tumours receive adjuvant endocrine treatment. Five years of aromatase inhibitor therapy produces a 40% reduction in 10-year mortality [1]. However, while the annual risk of mortality for ER-negative breast cancer decreases following the first five years after diagnosis, the annual rate remains constant for ER+ patients [2]. In fact, women with ER+ early-stage disease treated with 5 years of adjuvant endocrine therapy have a persistent risk of recurrence and death from breast cancer for at least 20 years after diagnosis [3]. Molecular studies have demonstrated that nodal and distant metastasis are highly similar to their matched primary tumours, implicating a continuation of original cancer [4-6]. However, the time between treatment and recurrence is often greater than that which can be explained by normal cell-doubling rates [7], implying cancer cells remain dormant in the body before re-awakening.

Residual dormant cancer cells are hypothesised to persist either by withdrawing from the cell cycle and transitioning to a quiescence state or by continuing to proliferate at a reduced rate, counter-balanced by cell death [8]. Reawakened dormant cells may become detectable after reaching a detection threshold or reactivated via increased angiogenesis, and/or escape from inhibitory microenvironment or immune effects [9, 10]. Dormancy is therefore considered a major mechanism underlying resistance to therapy, where dormant cells survive despite anti-proliferative endocrine treatment.

Resistance to endocrine therapy may occur at disease inception (*de novo* or innate resistance), but a larger proportion of patients acquire resistance during treatment (acquired/secondary resistance) [11]. Several mechanisms of endocrine resistance have been described previously [12, 13]. However, the majority of these findings are based on preclinical data obtained from cell lines and animal models. It is therefore difficult to know if these accurately reflect molecular changes in patient tumours.

Expression profiling of clinical samples, measuring the effect of, or predicting response to treatment has recently become feasible. However, experimental design issues, such as the difficulty in obtaining paired samples for comparison particularly for longer time intervals, makes it difficult to study changes within tumours [14]. For example, a previous study investigating tamoxifen failure compared samples from patients requiring salvage surgery with pre-treatment samples from an unrelated group of disease-free patients [15]. More recently, sequential patient-matched samples have been successfully utilised to determine treatment-induced dynamic changes in tumours at 2 weeks to 3 months, demonstrating the effectiveness of this approach [16-18].

For a variety of reasons, including being unfit for surgery, a proportion of patients receiving pre-surgical endocrine treatment do not have their tumours excised following 3-4 months of treatment. These long-term endocrine-treated tumours represent a unique group that can inform how tumours respond to extended oestrogen deprivation *in situ.* Having initially shrunk in size, some tumours remain at a steady volume and appear dormant, whilst others subsequently begin to regrow. We have utilised this unique cohort of sequential samples from patients receiving extended-neoadjuvant endocrine treatment to characterise luminal breast cancer dormancy and acquired resistance using as a novel clinical model.

## Methods

### Patients and samples

Breast cancer patients were treated with neoadjuvant letrozole (Femara, 2.5 mg; Novartis Pharma AG, Basel, Switzerland) for a minimum of four months, tumours were not removed either because patients declined or were unfit for surgery. The study was approved by the local regional ethics committee (07/S1103/26, August 2007) and all patients gave informed consent. Clinical characteristics of the tumours are given in Table 1. Cohort size with inclusion and exclusion criteria are given in Additional file 1: Figure S1. Patients with >40% initial decrease in tumour size by 4 months of treatment were included in the study. Those with no subsequent progression on imaging by the latest biopsy were classified as “dormant”, otherwise, they were classified as “acquired resistant” (Fig. 1a-b). For patients whose latest USS measurement was taken more than a month before surgery, changes in three widely used proliferation markers (MKI67, PCNA and, MCM2) were used to assist classification. Sequential tumour biopsies were taken with a 14-gauge needle before and after letrozole treatment and at the time of surgery. Fresh samples were snap-frozen in liquid nitrogen and each tumour sample confirmed to contain ≥ 50% cellularity and at least 60% tumour tissue using H&E sections. Following pulverisation of tissue with a membrane disruptor (Micro-Dismembrator U, Braun Biotech), phase separation was performed by guanidinium thiocyanate-phenol-chloroform extraction (Qiazol Lysis Reagent).

**Fig. 1.**
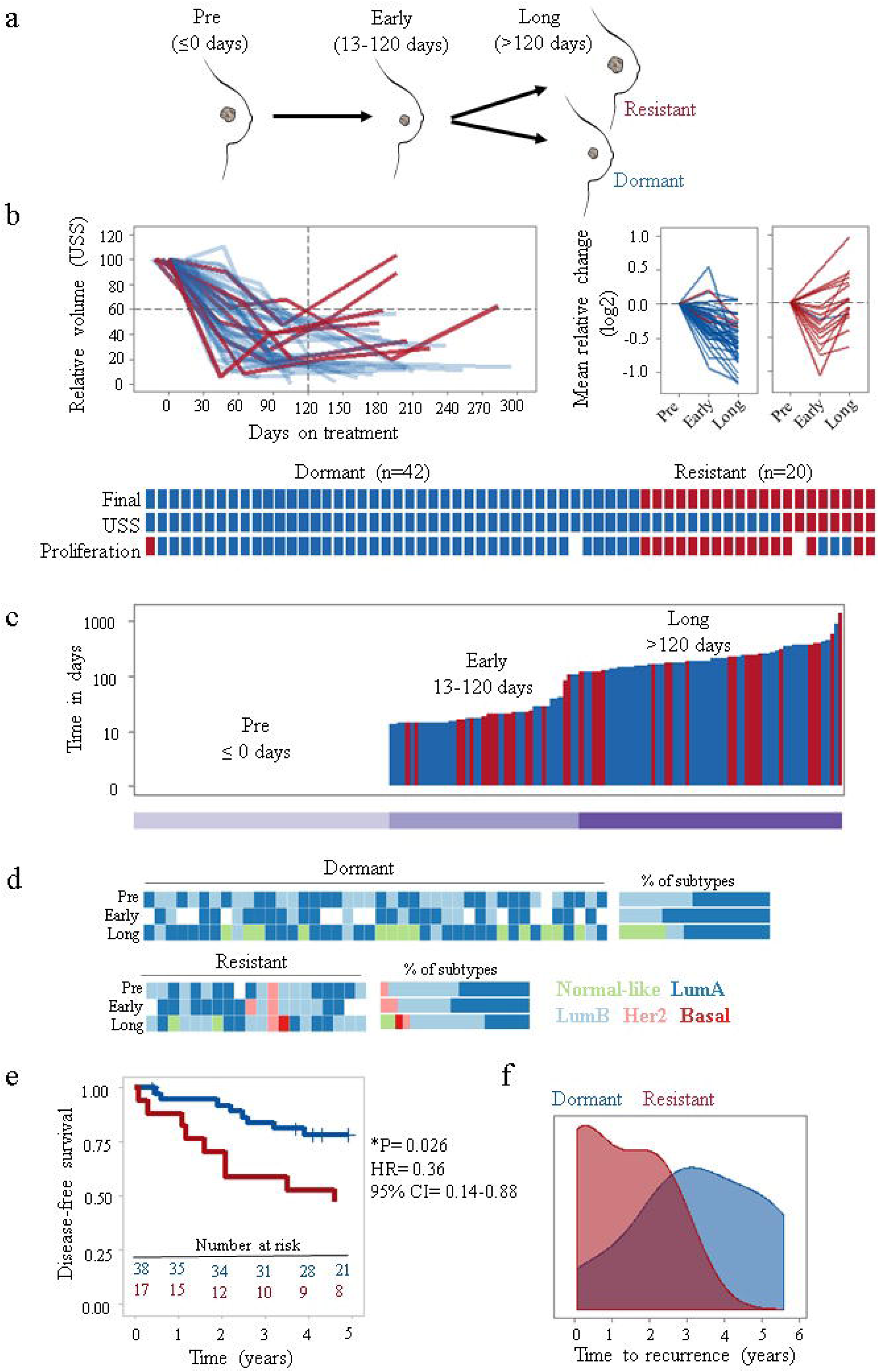
Long-term endocrine therapy as a clinical model to investigate breast cancer dormancy and acquired resistance. **a** Extended (4-45 months) letrozole treatment was exploited as a clinical model of breast cancer dormancy and acquired resistance. Sequential clinical samples from the same patient with no surgery and extended treatment were used to model clinical breast cancer dormancy and resistance. Before (pre, ≤0 days), early-on (early, 13-120 days) and long-term (long, >120 days) neoadjuvant letrozole treatment. **b** Dynamic change in tumour size by USS and mean expression of proliferation markers MKI67, PCNA and MCM2 were used to classify patients into two categories, dormant (blue) and resistant (red). Overall comparisons of classifications per patient based on USS and mean change in proliferation markers with final classification are shown. **c** The duration of letrozole treatment (days) for samples, each bar represents a sample. **d** Intrinsic subtype classification by PAM50 at each timepoint. **e** Kaplan-Meier plot showing overall survival probability in dormant vs resistant patients (log-rank test). **f** Density plot showing the distribution of time to recurrence in dormant and resistant patients.

**Table 1.** Patient characteristics.

|  | **Dormant**<br>n (%) | **Resistant**<br>n (%) | **Total**<br>n | **p value\*** |
| --- | --- | --- | --- | --- |
| *Total no of patients* | 42 | 20 | 62 |  |
| *Total no of samples* | 111 | 56 | 167 |  |
| *Age at diagnosis* |  |  |  |  |
| Mean | 75 | 72 |  |  |
| Range | 53-87 | 56-89 |  |  |
| *Tumour grade* |  |  |  | 0.39 |
| 1 | 6 (14.3) | 1 (5.0) | 7 |  |
| 2 | 27 (64.3) | 10 (50.0) | 37 |  |
| 3 | 8 (19.0) | 6 (30.0) | 14 |  |
| NA | 1 (2.4) | 3 (15.0) | 4 |  |
| *Tumour size* |  |  |  | 0.71 |
| T1 | 5 (11.9) | 4 (20.0) | 9 |  |
| T2 | 19 (45.2) | 9 (45.0) | 28 |  |
| T3 | 2 (4.8) | 2 (10.0) | 4 |  |
| T4 | 11 (26.2) | 4 (20.0) | 15 |  |
| NA | 5 (11.9) | 1 (5.0) | 6 |  |
| *Nodal status* |  |  |  | 0.36 |
| N0 | 27 (64.3) | 11 (55.0) | 38 |  |
| N1 | 8 (19.0) | 7 (35.0) | 15 |  |
| N2 | 1 (2.4) | 1 (5.0) | 2 |  |
| N3 | 1 (2.4) | 0 | 1 |  |
| NX | 1 (2.4) | 0 | 1 |  |
| NA | 4 (9.5) | 1 (5.0) | 5 |  |
| *Metastasis status* |  |  |  | 1.00 |
| M0 | 34 (80.9) | 18 (90.0) | 56 |  |
| M1 | 2 (4.8) | 0 | 2 |  |
| MX | 1 (2.4) | 1 (5.0) | 2 |  |
| NA | 5 (11.9) | 1 (5.0) | 6 |  |
| *ER score (Allred)* |  |  |  | 0.18 |
| 6 | 1 (2.4) | 0 | 1 |  |
| 7 | 6 (14.3) | 6 (30.0) | 12 |  |
| 8 | 35 (83.3) | 14 (70.0) | 49 |  |
| *HER status* |  |  |  | 0.69 |
| Negative | 35 (83.3) | 12 (60.0) | 47 |  |
| Positive | 6 (14.3) | 3 (15.0) | 9 |  |
| NA | 1 (2.4) | 5 (25.0) | 6 |  |
| *Molecular subtype<sup>#</sup>* |  |  |  | 1.00 |
| Luminal A | 21 (50.0) | 9 (45.0) | 30 |  |
| Luminal B | 20 (47.6) | 9 (45.0) | 29 |  |
| HER2 enriched | 0 | 1 (5.0) | 1 |  |
| Basal-like | 0 | 0 | 0 |  |
| Normal-like | 0 | 0 | 0 |  |
| NA | 1 (2.4) | 1 (5.0) | 2 |  |

2 \* Fisher exact test ( $p < 0.05$ two-tailed) <sup>#</sup> At diagnosis by PAM50 (genefu) NA: not available

### Gene expression profiling and analysis

RNA was extracted from the aqueous phase by column-based purification (RNeasy mini kit, Qiagen) and then labelled and hybridized (HumanHT-12 v4 Illumina BeadChip) according to manufacturer’s protocol (NuGEN) as previously described [19, 20]. Raw data was detection (*p* < 0.05, ≥ 3 samples) and quality filtered, log2 transformed, and quantile normalized using the Bioconductor lumi package [21]. Data is available from NCBI GEO under accession GSE111563. The analysis also includes data from 14 patients (42 samples, GSE59515) and 9 patients (24 samples, GSE55374) from previous studies [16, 19]. Hierarchical clustering analysis was performed using a complete linkage method. Pathway enrichment analysis and visualisation were performed using ReactomePA [22]. Differential gene expression analysis was performed with Rank Products [23]. The significance of differences was evaluated by using unpaired Wilcoxon test for two groups and ANOVA with post-hoc Tukey HSD for multiple comparisons.

### Proteomics analysis

Proteins were isolated from the organic phase of Qiazol [24]. Pellets were sonicated and dissolved in 1% SDS. Proteomics was performed using Thermo Q Exactive plus and Label-free Quantitation (LFQ). Peptides obtained from samples were analysed in mass spectrometry runs, serial samples from the patients were run on the same day. A modified version of Filter Aided Sample Preparation (FASP) was performed using serial digests with lysC and trypsin to generate two orthogonal fractions per sample [25, 26]. The mass spectrometry spectra generated in each run was used for relative quantitation of individual peptides. Normalization and quantifications of peptides were performed using MaxLFQ and MaxQuant [27]. A total of 6251 protein groups were identified. Data was log2 transformed and missing values were imputed as the minimum observed value in each sample. The data have been deposited to the ProteomeXchange Consortium via the PRIDE [28] partner repository with the dataset identifier PXD009328.

### Immunohistochemistry and scoring

Formalin-fixed paraffin-embedded (FFPE) sections were processed using an automated stainer (Leica Biosystems, Bond III). Heat-induced epitope retrieval for both antibodies was done by 30-minute incubation in citrate based pH 6.0 epitope retrieval (ER1) solution followed by incubation in 3.5 N HCl for 15 min at room temperature as suggested by Haffner *et.al.* [29]. For 5-methylcytosine (5-mC) and 5-hydroxymethylcytosine (5-hmC) detection, mouse monoclonal 5 methylcytosine specific [33D3] (Abcam, ab 10805) and rabbit polyclonal 5-hydroxylmethylcytosine (Active Motif, 39769) antibodies were used respectively. Both antibodies were used at 1/1000 dilution and were incubated for 15 min. Detection was performed by using secondary antibody-horseradish peroxidase (HRP) conjugates and substrate-chromogen (DAB). After staining, slides were counterstained with haematoxylin. Nuclear staining in epithelial cells was evaluated using an H-score obtained by multiplying the intensity of the stain (0: no staining; 1: weak staining; 2: moderate staining; 3: intense staining) by the percentage of cells (H-score range, 0 to 300).

## Results

### Long-term endocrine therapy as a model of dormancy and acquired resistance

A cohort of 62 primary breast cancer patients receiving at least 4 months of endocrine treatment (Fig. 1a) were stratified into two groups, ‘dormant’ and ‘acquired resistant’ based on dynamic changes in tumour size and proliferation (Fig. 1b). Patient-matched sequential samples were available at three time-points, before (pre, ≤0 days), early-on (early, 13-120 days) and long-term (long, >120 days) treatment. Dormant and acquired resistant samples were distributed uniformly with respect to time on treatment, and duration at each time point was not significantly different between response groups (Table 1). For long-term treatment, the mean and range were 186 (121-884) days and 226 (121-1366) days, for dormant and acquired resistant patients respectively (Fig. 1c).

There were no significant differences in patient clinico-pathological features between response classes before treatment (Table 1). However, PAM50 intrinsic molecular subtypes were found to change during endocrine treatment (Fig. 1d). These changes were consistent with known associations with outcome, with all dormant tumours either remaining the same, or switching to better prognosis luminal A or normal-like tumours. For resistant tumours, however, 25% (5 out of 20) switched to a subtype of worse prognosis (Fig. 1d).

As expected, Kaplan-Meier survival analysis demonstrated significantly worse outcomes for resistant compared to dormant patients (log rank, *p* = 0.026, Fig. 1e). Recurrence rates for dormant and resistant patients were 21% (9/42) and 45% (9/20), respectively. Moreover, resistant patients suffered significantly earlier recurrences compared to dormant patients (*p* = 0.05; range = 26-947 vs 136-2042 days; Fig. 1f).

### Distinct transcriptomic changes under long-term letrozole treatment

Unsupervised analysis was performed to consider whether sequential samples displayed greater similarity between response classes or treatment duration. Hierarchical clustering using the 500 genes with the highest variance across all samples revealed two main subclasses, seemingly driven by time on treatment, with resistant and dormant tumours indistinguishable. When long-term treated samples were considered alone, two clusters did emerge, the larger of which contained a majority of dormant samples (79%), whilst the second had a roughly even proportion of dormant (48%) and resistant (52%) samples (Fig. 2a). Similarly, a multidimensional scaling (MDS) plot revealed consistent changes over time in response to treatment for both dormant and acquired resistant samples (Fig. 2b), although long-term dormant samples were much more distinguishable from pre-treatment samples than the long-term acquired resistant samples (Fig. 2b).

**Fig. 2.**
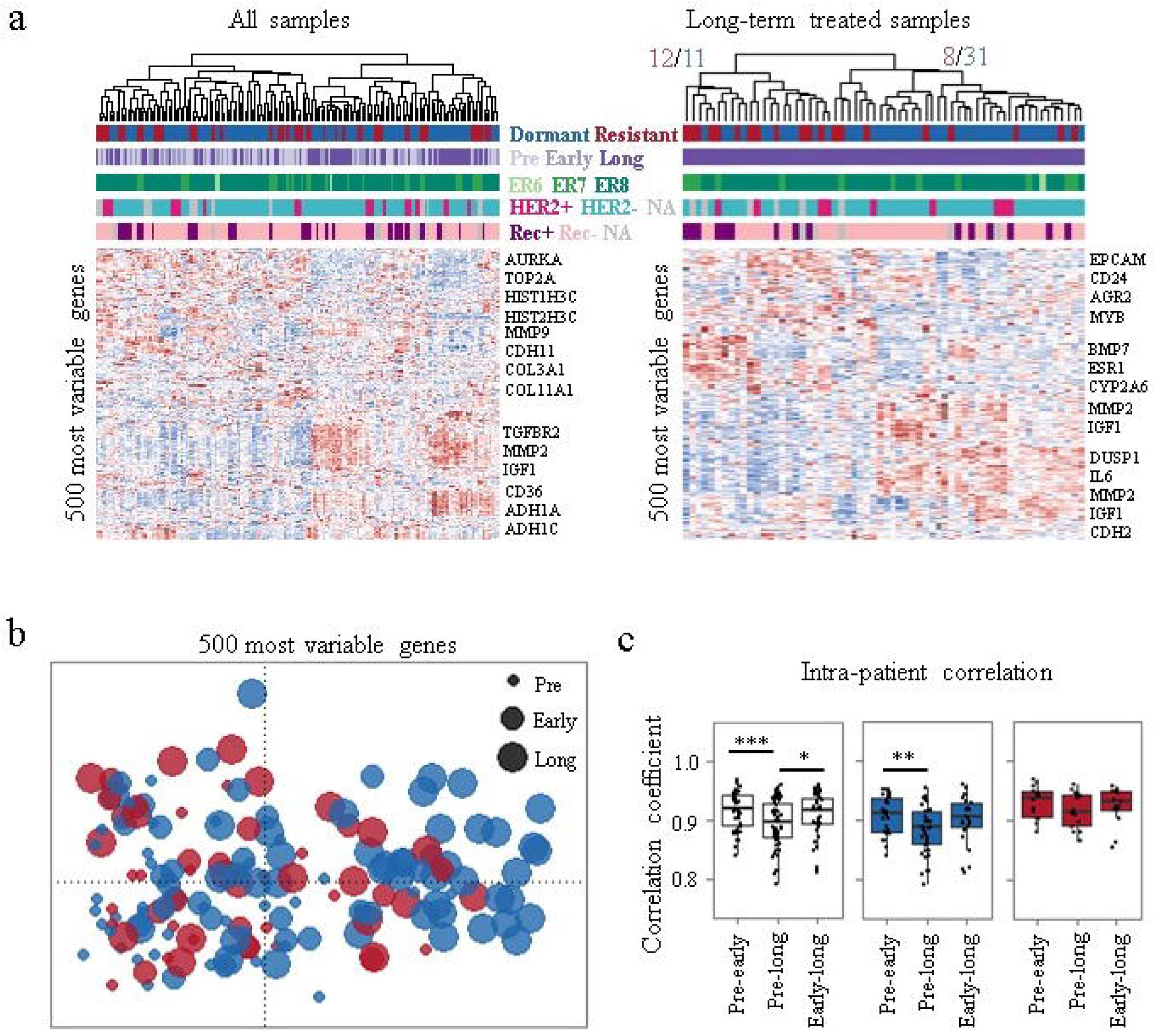
Distinct transcriptomic changes during long-term aromatase inhibitor treatment. **a** Unsupervised hierarchical clustering with most variant 500 features. **b** Multidimensional scaling (MDS) plot using most variant 500 genes across all timepoints. Each dot corresponds to a sample and sizes represent the duration of treatment. **c** Intra-patient (comparison of samples from the same patient) correlations are shown. Dormant (blue); resistant (red); Before (pre, ≤0 days), early-on (early, 13-120 days) and long-term (long, >120 days) neoadjuvant letrozole treatment; ^∗∗∗^ *p* < 0.001; ^∗∗^ *p* < 0.01; ^∗^ *p* < 0.05.

Correlations between tumours from different individuals (inter-patient) remained similar at each time point and were not different between response classes (not shown). However, correlations between matched sequential samples (intra-patient) revealed that pre-treatment samples were significantly (*p* = 0.01) less similar to their long-term treated pairs (median = 0.89, range = 0.74-0.95) than their early-on treatment pairs (median = 0.91, range = 0.84-0.95) (Fig. 2c). However, when divided by dormancy status this finding was only significant (*p* = 0.01) for dormant patients (Fig. 2c), suggesting that dormant tumours continue to diverge transcriptionally, whereas acquired resistant tumours do not consistently differ after initial or extended treatment, as mirrored in the MDS representation (Fig. 2b).

### Changes in genes/pathways following long-term letrozole treatment

Pairwise Rank Product analysis (pre- vs. long-term treatment, FDR < 0.01) of dormant patients identified 2319 genes significantly differentially expressed (1063 down- and 1256 up-regulated) (Additional file 2: Table S1). These genes were significantly enriched (*p* < 0.01) for a total of 62 and 26 pathways, respectively (Additional file 2: Table S2), including reductions in cell cycle, senescence, DNA methylation and an increase in extracellular matrix (ECM) organization. These findings are consistent with previous studies of patient-matched sequential endocrine treated samples [16-18]. Acquired resistant tumours displayed much fewer consistently differentially expressed genes (238, 63 down- and 175 up-regulated) between long-term treated and pre-treatment samples (Additional file 2: Table S3). Genes that were up-regulated in resistant patients were enriched for several of the same pathways as dormant tumours (ECM organization, Elastic fibre formation and Platelet degranulation), but down-regulated genes were much more variable (Additional file 2: Table S4; Fig. 3a-b).

**Fig. 3.**
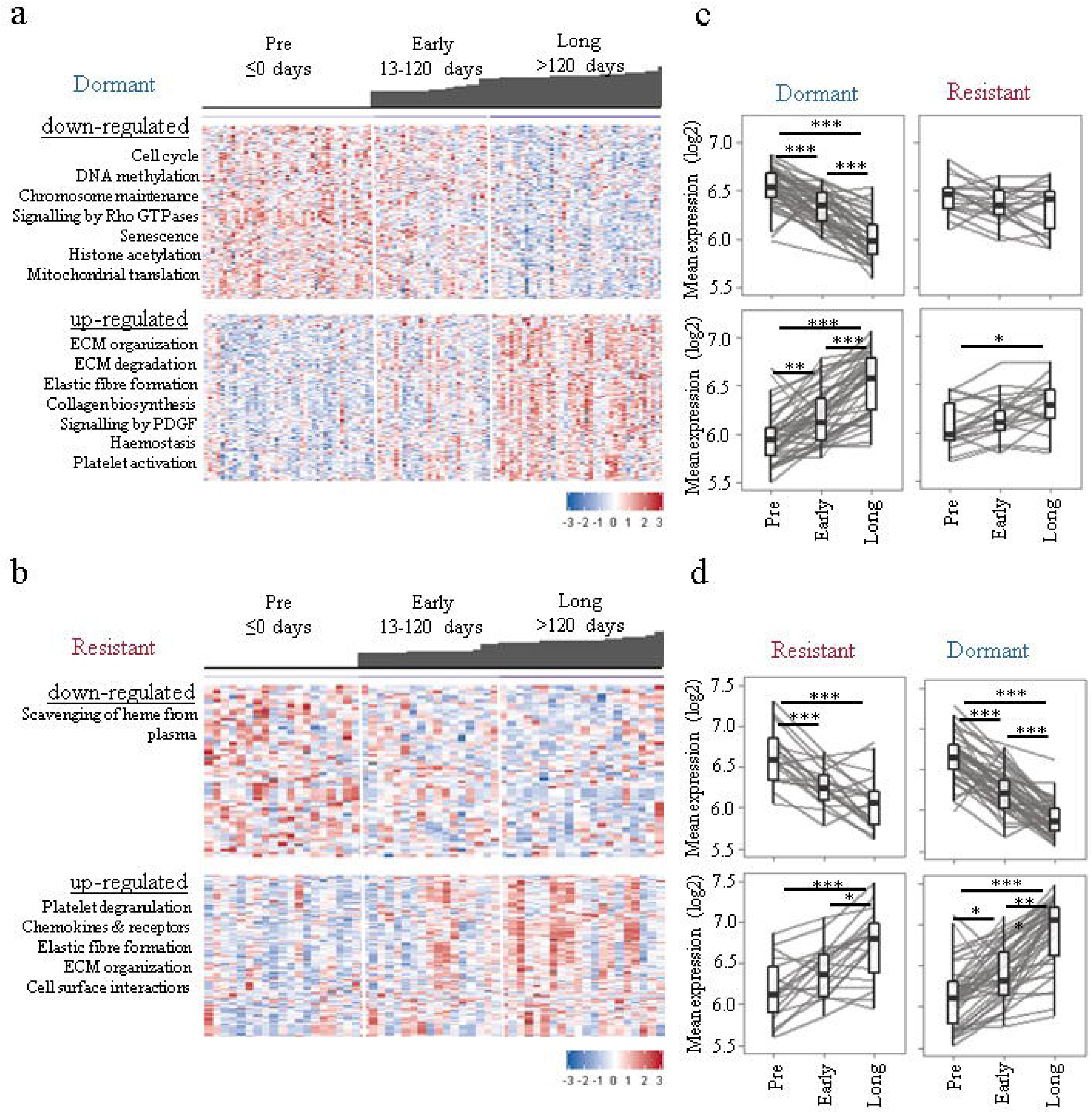
Long-term endocrine treatment is associated with cell cycle, senescence, epigenetic regulation and ECM-associated pathways. Differentially expressed genes between pretreatment and long-term treated samples of dormant (**a**) and resistant (**b**) patients were determined. Heat-maps showing change in down- and up-regulated genes’ expression in dormant (**a**) and resistant (**b**) samples. Each column represents a sample and each row a gene. Colours are log2 mean-centred values with red indicating high values and blue indicating low expression. Bar plots on top of heat-maps represent the time on treatment (days) for each sample. **c-d** Graphs show dynamic changes in mean expression of differentially expressed genes in response classes. ^∗∗∗^*p* < 0.001; ^∗∗^ *p* < 0.01; ^∗^*p* < 0.05.

Having determined that dormant and acquired resistant tumours have somewhat distinct changes during treatment at the molecular level, the question remained as to whether these changes tend to occur at earlier time points or were specific to long-term treatment. For dormant tumours, differential expression begins early-on, but becomes more pronounced at later timepoints (Fig. 3a). Down-regulated genes were most evident early-on treatment for resistant patients, consistent with their initial response to treatment; whilst up-regulated genes were most changed after long-term treatment, potentially suggesting that these genes may mediate acquired resistance (Fig. 3b). We further examined whether differentially expressed genes identified in each response class were shared (Fig. 3c-d). Both down- and up-regulated genes identified in resistant tumours were significantly changed (*p* < 0.01) in dormant patients (Fig. 3d). However, only up-regulated genes identified in dormant patients were significantly up-regulated in resistant patients without any change in down-regulated genes (Fig. 3c), implicating a partial lack of response to treatment at the molecular level in acquired resistance patients.

### A potential role of epigenetic regulation in acquired resistance

The above findings suggested that therapy-induced dynamic changes in genes and pathways are common features of long-term treatment, rather than being specific to dormant or resistant phenotypes. This led us to perform comparative analysis of dormant and acquired resistant tumours at long-term time-point to identify any specific differences. Unpaired Rank Product analysis (FDR < 0.01) revealed a total of 419 genes (170 down- and 249 up-regulated) to be differentially expressed between long-term treated dormant and resistant tumours (Additional file 2: Table S5; Fig. 4a). These genes were significantly enriched in 27 pathways (*p* < 0.05), including several epigenetics-related pathways, including “DNA methylation”, “PRC2 methylates histones and DNA”, “histone acetyl transferases (HATs) acetylate histones”, “epigenetic regulation of gene expression” as well as senescence and cell cycle (Additional file 2: Table S6; Fig. 4b). Examination of the expression of these genes alone, demonstrated that they could partially separate dormant from majority of resistant tumours (Fig. 4c). Single-sample Gene Set Enrichment Analysis (ssGSEA) [30] was performed to quantitatively score the activity of differentially expressed genes in every sample. The differentially up-regulated genes’ score was significantly higher in acquired resistant compared to dormant tumours under early-on (*p* < 0.05) as well as long-term (*p* < 0.001) treatment (Fig. 4d).

**Fig. 4.**
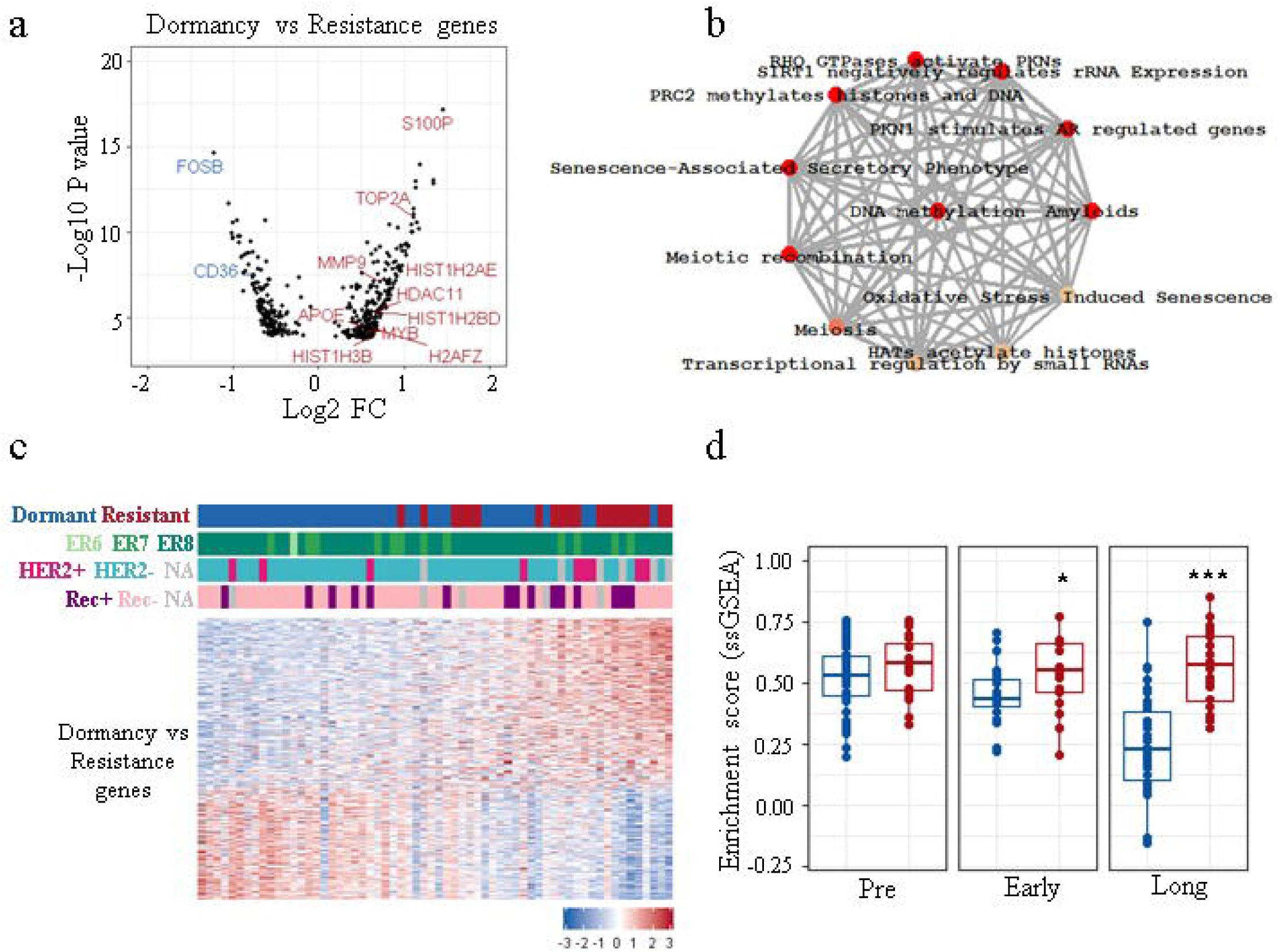
Comparative analysis of dormant and resistant tumours. **a** Volcano plot showing differentially expressed genes between long-term treated dormant and resistant tumours (dormancy vs resistance genes). Some up- and down-regulated genes in resistant tumours are highlighted in red and blue, respectively. **b** Significantly enriched pathways for dormancy vs resistance genes (*p* < 0.01; ReactomePA). **c** Heatmap showing partial separation of long-term treated dormant and resistant samples using dormancy vs resistance genes. Colours are log2 mean-centred values with red indicating high and blue indicating low expression. Genes are sorted by FC values from most to least up/down-regulated. Samples are sorted by sum expression of up-regulated genes. **d** Comparison of ssGSEA scores of dormancy vs resistance up-regulated genes between dormant and acquired resistant tumours. Dormant (blue); resistant (red); Before (pre, ≤0 days), early-on (early, 13-120 days) and long-term (long, >120 days) neoadjuvant letrozole treatment; ^∗∗∗^ *p* < 0.001; ^∗^*p* < 0.05.

Our results prompted us to examine whether the changes we observed in clinical samples were similarly changed in experimental models of resistant breast cancer cells. Oestrogen receptor-positive MCF7 cells stably transfected with the aromatase gene (MCF7aro cells) and long-term oestrogen-deprived (LTED) breast cancer cells have been widely used to understand mechanisms of aromatase inhibitor resistance *in vitro.* Examining two publicly available gene expression datasets (GSE10879 and GSE10911) demonstrated that genes differentially expressed between acquired resistant and dormant tumours were significantly enriched in aromatase inhibitor-resistant cells compared to sensitive/control cells (Fig. 5a). In two out of three *in vitro* studies with dynamic gene expression data from LTED MCF7 cells with an initial decrease in ssGSEA scores mimicking the dormancy/responsive state was followed by a later-on increase representing acquired resistance (Fig. 5b), further validating our results and emphasizing the utility of these *in vitro* models. Interestingly, no significant difference was observed in tamoxifen- and fulvestrant-resistant MCF7 cells compared to drug-sensitive control cells (Fig. 5c) suggesting the specificity of the results to aromatase inhibitor therapy resistance.

**Fig. 5.**
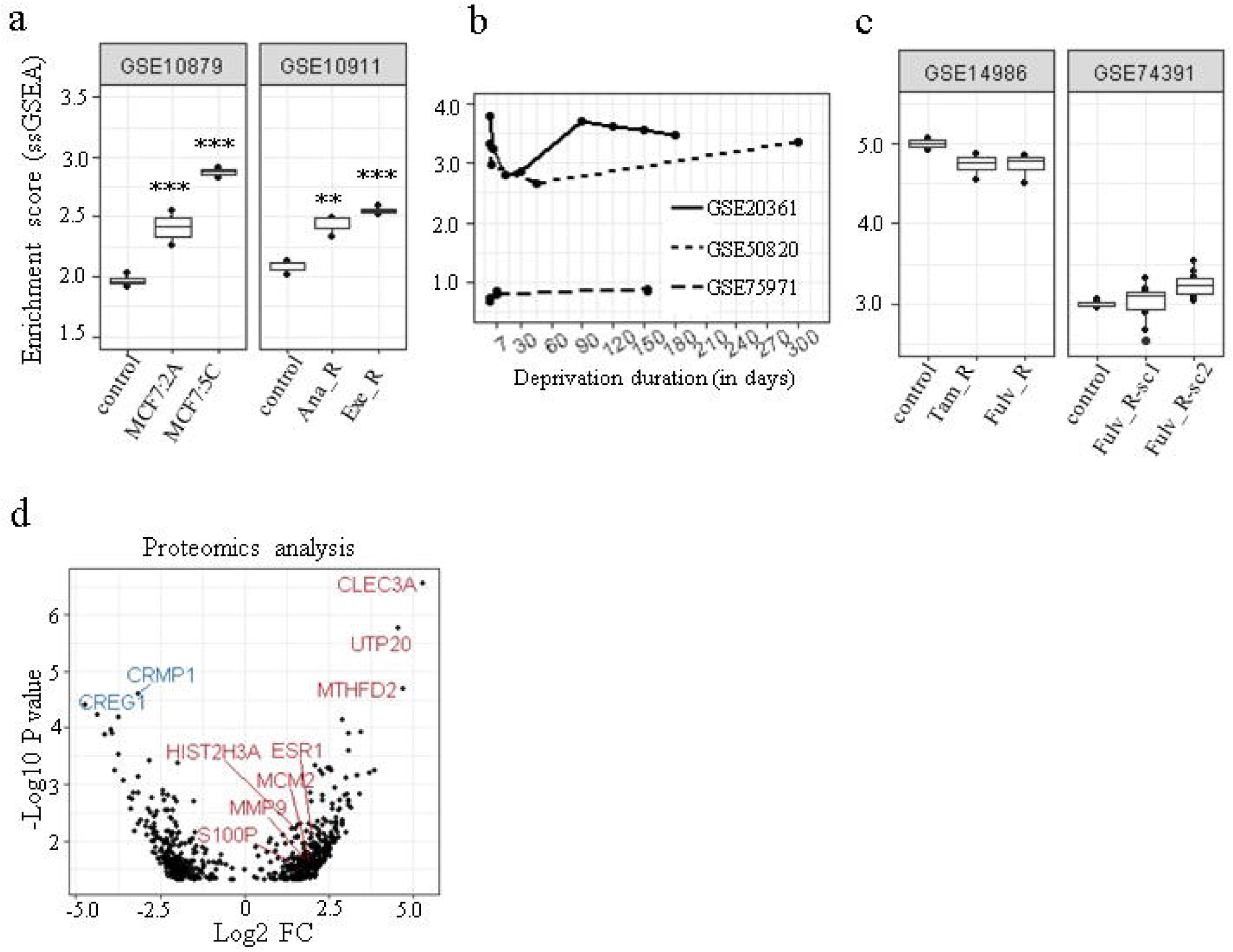
Validation of results using *in vitro* gene expression data from resistant cell lines and proteomics analysis. **a** Normalised enrichment scores of differently up-regulated genes calculated using single sample gene set enrichment analysis (ssGSEA) in aromatase inhibitor-resistant cells. Scores were significantly higher (^∗∗^ *p* < 0.01, ^∗∗∗^ *p* < 0.001) in two aromatase inhibitor-resistant cell lines, MCF7:2A and MCF7:5C, which were clonally derived from MCF7 breast cancer cells following long-term estrogen deprivation (LTED) compared to control/sensitive MCF7 cells (n = 4). Anastrozole-resistant (Ana_R) and exemestane-resistant (Exe_R) MCF7aro cells had significantly higher scores compared to control (n = 3). **b** Dynamic changes in enrichment scores of LTED MCF7 cells in three different datasets. **c** Scores in tamoxifen-resistant (Tam_R) and fulvestrant-resistant (Fulv_R) and drug-sensitive (control) MCF7 cells (n = 4, n = 10; sc: subclone). **d** Volcano plot showing differentially expressed proteins between long-term treated dormant and resistant tumours (*p* < 0.05). Some overlapping features between transcriptomics and proteomics analysis and the most up- and down-proteins are highlighted in red and blue, respectively

In addition, proteomic analysis of a subset of samples was performed which revealed differential expression in 656 proteins (279 down-, 377 up-regulated) between long-term treated dormant and resistant tumours (Rank Product; *p* < 0.05; n = 10; Additional file 2: Table S7; Fig. 5d). A total of 36 features including S100P and HIST2H3A (H3.2) overlapped between proteomics and transcriptomics, validating the results with a different approach.

Next, we considered which drugs might be able to reverse the differences in gene expression between dormant and resistant tumours using Connectivity Map (CMap), the pattern matching software, and a collection of gene-expression profiles from cultured human cells treated with various treatments [31]. Drugs that are negatively associated with the resistance signature could potentially reverse the differences observed. A histone deacetylase (HDAC) inhibitor Trichostatin A had the second lowest score (−0.90) and has previously been shown to re-sensitise tamoxifen-unresponsive ER-negative breast cancer cells *in vitro* [32]. Clioquinol an anti-parasitic and apoptotic drug with HDAC inhibitory effects [33] was also highlighted. Letrozole had a positive score of 0.89 further confirming the reliably of the predictions and hypothetical scores calculated by CMap. Furthermore, differentially expressed genes were uploaded to Enricher (ENCODE Histone modification 2015 dataset) [34] to determine histone modification enrichment. Two H3 lysine methylation modifications (H3K27me3 and H3K4me1) were enriched significantly (Adjusted *p* = 0.0003 and *p* = 0.004, respectively) whereas no enrichment for histone acetylation was determined.

Finally, immunohistochemical evaluation of FFPE sections revealed significantly lower global 5-mC and 5-hmC levels in resistant tumours compared to dormant tumours under extended treatment (Fig. 6a-b). Significantly lower 5-hmC levels in acquired resistant compared to dormant tumours were also observed at early on-treatment (Fig. 6b), suggesting hypomethylation may be predictive of emergence from dormancy.

**Fig. 6.**
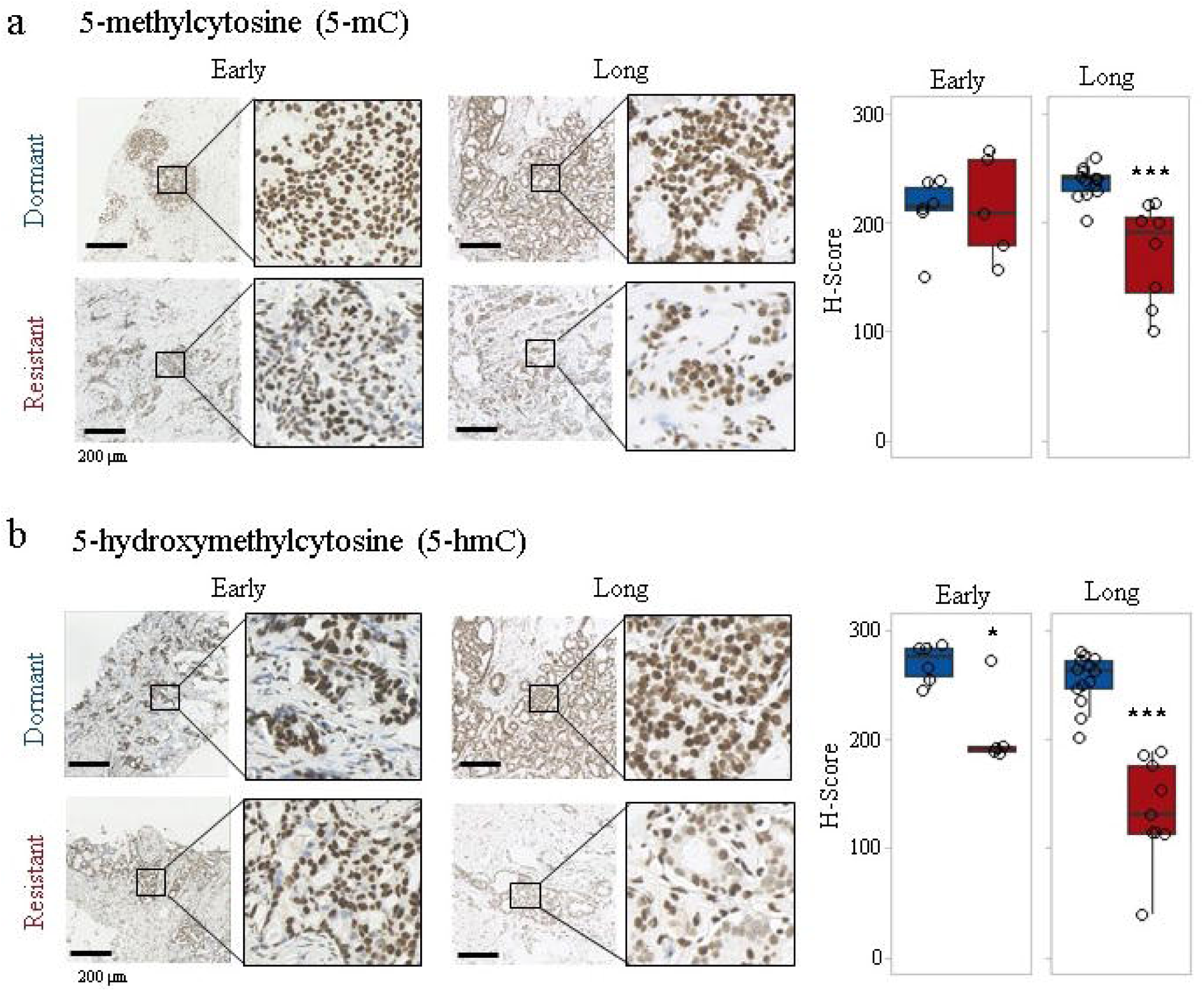
Immunohistochemical evaluation of **a** 5-methylcytosine (5-mC) and **b** 5-hydroxymethylcytosine (5-hmC) in FFPE sections from letrozole-treated samples. Representative images in dormant and resistant tumours are shown. Boxplots show distributions of semi-quantitative intensity scores of 5-mC (^∗^ *p* < 0.05, ^∗∗∗^ *p* < 0.001, n = 512) and 5-hmC (^∗∗∗^ *p* < 0.001, n = 5-13) levels in dormant and resistant patients. Early-on (early, 13-120 days) and long-term (long, >120 days) neoadjuvant letrozole treatment.

## Discussion

Understanding the mechanisms underlying the maintenance of and escape from dormancy have great importance considering most cancer-related deaths are caused by metastasis rather than the primary tumour. In this study, we describe the first sequential patient-matched clinical dataset of extended endocrine treatment in breast cancer. The results highlight the difficulty of distinguishing dormant and resistant tumours, with dynamic molecular changes of treatment being highly similar between the groups. However, comparative analysis revealed a set of genes significantly up-regulated in resistant patients within the first months of letrozole treatment suggesting a predictive role for changes in DNA methylation.

Failure to reduce proliferation after 2 weeks of endocrine treatment [16, 35] may well identify patients that are innately resistant; however, acquired resistance remains a greater challenge in terms of identifying biomarkers and appropriate alternative or combination therapies [36]. Many of the transcriptomic changes identified in long-term treated dormant tumours are shared by some, but not all resistant tumours, providing further evidence of resistance heterogeneity [37] where dormant tumours share similar molecular changes, but there are a variety of escape mechanisms that lead to acquired resistance.

In the present study, paired differential expression analysis demonstrated that dormant tumours continue to change under long-term treatment. Some of the identified dormancy-related pathways such as cell cycle arrest and senescence have established roles in metastasis dormancy [38] further supporting the relevance of our clinical model, with senescence-associated secretory phenotype (SASP) recently suggested to regulate breast cancer dormancy and relapse [39]. As in short-term responsive tumours [16], ECM organization and degradation were significantly up-regulated in dormant tumours. ECM remodelling and its degradation by matrix metalloproteases (MMP) have previously been suggested to regulate the switch between dormancy and metastatic growth [40].

The most up-regulated gene in resistant tumours *S100P*, previously shown to be an inducer of breast cancer metastasis correlated with decreased survival [41]. Recently, *S100P* hypomethylation in blood was demonstrated to be inversely correlated with tissue *S100P* expression and significantly associated with breast cancer, implicating *S100P* as a potential diagnostic marker [42]. High plasma *S100P* levels have also been correlated with poor prognosis in metastatic breast cancer patients, with levels decreasing following treatment, suggesting a role of S100P in dynamic monitoring of response [43]. In the present study, S100P gene expression and protein levels were significantly higher in resistant tumours after long-term treatment, as well as being differentially expressed before treatment supporting its potential role as a therapeutic target [44] and a predictive marker.

Comparative analysis of dormant and resistant samples after extended treatment revealed a set of genes enriched for DNA methylation and histone acetylation/deacetylation, indicating the involvement of epigenetic regulation in escape from dormancy. Epigenetic alterations are recognized to occur in breast cancer. DNA methyltransferase (DNMT) and HDAC inhibitors have been shown to exert encouraging effects in the disease [45]. Recently, the potential role of epigenetic changes in regulating dormancy and reactivation state has been suggested to explain the reversible (on/off) nature of dormancy [46].

Breast cancer “CpG island methylator phenotype” (CIMP), revealed by genome-wide methylation analysis of metastatic breast cancers where a large number of genes are hypermethylated, and has been suggested to be informative for metastatic potential [47]. A significant correlation between pre-treatment global DNA methylation with neoadjuvant chemotherapy response in rectal cancer has been reported [48]. Though DNA hypomethylation was the first epigenetic alteration identified in cancer, its molecular process and effects are not well understood yet [49]. In addition, 5-hmC levels were shown to correlate with differentiation status, with higher levels in more differentiated [29]. In addition, alterations in DNA methylation in LTED MCF7 cells have been previously reported [50]. Our results provide evidence for loss of global DNA methylation process in resistant tumours and strengthen the case to use these models for further study. The global decrease in 5-mC may account for the observed reduction in 5-hmC levels, since 5-mC is converted to 5-hmC. On the other hand, at early-on time point, 5-hmC levels were significantly reduced with no significant change in 5-mC levels suggesting an independent role of 5-hmC mark. Hypomethylated cancer cells have been suggested to be selected to form tumours with increased malignancy [49]. We suggest that hypomethylation in resistant tumours may reflect dedifferentiation process inducing stem-cell-like cell formation. Determining the time point at which that hypomethylation starts, which would allow intervening before it starts to prevent therapy resistance, needs further investigation.

The main genes significantly enriched for epigenetics-associated pathways in the present study are core histone (H3, H4, H2B) genes. Well-known epigenetics-associated genes such as DNA methyltransferase (DNMT) were not differentially expressed in the present study. Therefore, it might be suggested that observed changes in histone gene levels may simply reflect the high proliferation rate in resistant tumours. Although these histone genes are replication-dependent and their levels increase during DNA replication [51], we suggest that elevated levels may also result from disrupted DNA replication and packaging. Deregulation of histone H2A and H2B were associated with anthracycline resistance in breast cancer cells and reversed by HDAC small molecule inhibitors [52]. Furthermore, up-regulation of replication-dependent core histone proteins has been suggested to be selective indicator of ER-mediated MCF7 cell proliferation regardless of proliferation-rate [53]. Also, observed global loss of DNA methylation in resistant tumours suggests dynamic regulation of gene transcription under letrozole therapy. Therefore, histone up-regulation and alterations in epigenetic pathways observed in our study may mediate reactivation of ER signalling in resistant tumours, rather than simply mirroring the degree of proliferation.

Our results indicate alterations both in DNA methylation, and histone modifications suggesting a cooperative interplay between them to mediate acquired resistance. HDAC inhibitors, which have been shown to regulate DNA methylation [54], may be successful clinically as second-line drugs alone or in combination following endocrine therapy failure as there is growing evidence for their tumour selective action [55, 56]. A time-dependent role for HDACs in leukaemia has been shown [57] and may also be critical in determining when to start HDAC inhibition therapy to successfully treat endocrine resistant patients. Whether or not the epigenetic alterations are triggers of reawakening and if the timely use of epigenetic drugs can prevent acquired resistance warrants further investigation.

## Conclusions

We have performed the first study of sequential tumour samples from breast cancer patients receiving extended neoadjuvant endocrine treatment as a clinical model of dormancy and acquired resistance. Our analysis suggests that molecular differences between dormant and resistant tumours are initially subtle, becoming more obvious only after extended treatment. This study emphasizes alterations in DNA methylation in the first months of treatment may predict which patients will eventually develop acquired resistance. We provide valuable evidence that epigenetic drugs such as HDAC inhibitors may have a role in treating endocrine resistance.

## Abbreviations

5-hmC: 5-hydroxymethylcytosine
5-mC: 5-methylcytosine
CIMP: CpG island methylator phenotype
cMap: Connectivity map
DAB: 3,3′-diaminobenzidine
DNMT: DNA methyltransferase
ECM: extracellular matrix
ER: oestrogen receptor
FASP: Filter Aided Sample Preparation
FDR: false discovery rate
FFPE: formalin-fixed paraffin embedded
HATs: histone acetyl transferases
HCl: hydrochloric acid
HDAC: histone deacetylase
HRP: horseradish peroxidase
LFQ: Label-free Quantitation
LTED: long-term oestrogen deprivated
MDS: multidimensional scaling
MMP: matrix metalloproteases
SASP: senescence-associated secretory phenotype
ssGSEA: Single-sample Gene Set Enrichment Analysis

## Declarations

### Ethics approval and consent to participate

All patients provided informed consent and sample collection was approved by the local research ethics committee (Lothian Local Research Ethics Committee 03, REC Reference number 07/S1103/26, approval date 13/08/2007).

### Consent for publication

Not applicable

### Availability of data and material

The microarray dataset generated during the current study is available in NCBI GEO under accession GSE111563 [https://www.ncbi.nlm.nih.gov/geo/query/acc.cgi?acc=GSE111563]. The analysis also includes previously published microarray data under accession numbers GSE59515 [https://www.ncbi.nlm.nih.gov/geo/query/acc.cgi?acc=GSE59515], and GSE55374 [https://www.ncbi.nlm.nih.gov/geo/query/acc.cgi?acc=GSE55374]. The proteomics dataset generated during the current study is available in PRIDE with the identifier PXD009328 [https://www.ebi.ac.uk/pride/archive/]. Publicly available resistant cell line gene expression datasets GSE10879 [https://www.ncbi.nlm.nih.gov/geo/query/acc.cgi?acc=GSE10879], GSE10911 [58], GSE20361 [59], GSE50820 [60], GSE75971 [61], GSE14986 [62], GSE74391 [63] were also analysed.

## Authors’ contributions

AT, JMD and AHS conceived the study. CS and AT generated the transcriptome dataset. JW conducted proteomics and supported proteomics data analysis. DP, AL, and AHS provided help with the data analysis. AF and LR provided technical support with tissue collection and processing. LR, JST, and JMD co-ordinated the collection and assessment of clinical samples. CS analysed and interpreted the data and drafted the manuscript. AHS supervised the project and helped to write and edit the manuscript. All authors read and approved the final manuscript.

## Acknowledgements

Not applicable

## Authors’ information

Not applicable

## Additional files

**Additional file 1: FigureS1.** Consort diagram showing the cohort and sample sizes. Patient and sample sizes in each group are shown with inclusion and exclusion criteria. (JPG 81 kb)

**Additional file 2: TableS1-S7.** Tables for lists of differentially expressed genes and proteins, and enriched pathways. **Table S1**. Differentially expressed genes between long-term treated and pre-treatment samples of dormant patients. **Table S2.** Enriched pathways for differentially expressed genes between long-term treated and pre-treatment samples of dormant patients. **Table S3.** Differentially expressed genes between long-term treated and pre-treatment samples of acquired resistant patients. **Table S4.** Enriched pathways for differentially expressed genes between long-term treated and pre-treatment samples of acquired resistant patients. **Table S5.** Differentially expressed genes between long-term treated dormant and resistant tumours. **Table S6.** Enriched pathways for differentially expressed genes between long-term treated dormant and resistant tumours. **Table S7.** Differentially expressed proteins between long-term treated dormant and resistant tumours. (XLSX 283 kb)

## References

1. EBCTCG. Aromatase inhibitors versus tamoxifen in early breast cancer: patient-level meta-analysis of the randomised trials. Lancet. 2015;386(10001):1341−1352.

2. Demicheli R, Ardoino I, Boracchi P, Coradini D, Agresti R, Ferraris C, et al. Recurrence and mortality according to estrogen receptor status for breast cancer patients undergoing conservative surgery. Ipsilateral breast tumour recurrence dynamics provides clues for tumour biology within the residual breast. BMC Cancer. 2010;10:656.

3. Pan HC, Gray R, Braybrooke J, Davies C, Taylor C, McGale P, et al. 20-Year Risks of Breast-Cancer Recurrence after Stopping Endocrine Therapy at 5 Years. N Engl J Med. 2017;377(19):1836−1846.

4. Weigelt B, Glas AM, Wessels LF, Witteveen AT, Peterse JL, van’t Veer LJ. Gene expression profiles of primary breast tumors maintained in distant metastases. Proc Natl Acad Sci U S A. 2003;100(26):15901−15905.

5. Tang MH, Dahlgren M, Brueffer C, Tjitrowirjo T, Winter C, Chen Y, et al. Remarkable similarities of chromosomal rearrangements between primary human breast cancers and matched distant metastases as revealed by whole-genome sequencing. Oncotarget. 2015;6(35):37169−37184.

6. Kroigard AB, Larsen MJ, Thomassen M, Kruse TA. Molecular Concordance Between Primary Breast Cancer and Matched Metastases. Breast J. 2016;22(4):420−430.

7. Demicheli R, Terenziani M, Bonadonna G. Estimate of tumor growth time for breast cancer local recurrences: rapid growth after wake-up? Breast Cancer Res Treat. 1998;51(2):133−137.

8. Uhr JW, Pantel K. Controversies in clinical cancer dormancy. Proc Natl Acad Sci U S A. 2011;108(30):12396−12400.

9. Sosa MS, Bragado P, Aguirre-Ghiso JA. Mechanisms of disseminated cancer cell dormancy: an awakening field. Nat Rev Cancer. 2014;14(9):611−622.

10. Dittmer J. Mechanisms governing metastatic dormancy in breast cancer. Semin Cancer Biol. 2017;44:72−82.

11. Selli C, Dixon JM, Sims AH. Accurate prediction of response to endocrine therapy in breast cancer patients: current and future biomarkers. Breast Cancer Res. 2016;18(1):118.

12. Clarke R, Tyson JJ, Dixon JM. Endocrine resistance in breast cancer‐‐An overview and update. Mol Cell Endocrinol. 2015;418 Pt 3:220−234.

13. Ma CX, Reinert T, Chmielewska I, Ellis MJ. Mechanisms of aromatase inhibitor resistance. Nature Reviews Cancer. 2015;15(5):261−275.

14. Sims AH, Bartlett JMS. Approaches towards expression profiling the response to treatment. Breast Cancer Res. 2008;10(6).

15. Vendrell JA, Robertson KE, Ravel P, Bray SE, Bajard A, Purdie CA, et al. A candidate molecular signature associated with tamoxifen failure in primary breast cancer. Breast Cancer Res. 2008;10(5):R88.

16. Turnbull AK, Arthur LM, Renshaw L, Larionov AA, Kay C, Dunbier AK, et al. Accurate Prediction and Validation of Response to Endocrine Therapy in Breast Cancer. J Clin Oncol. 2015;33(20):2270−2278.

17. Dunbier AK, Ghazoui Z, Anderson H, Salter J, Nerurkar A, Osin P, et al. Molecular Profiling of Aromatase Inhibitor-Treated Postmenopausal Breast Tumors Identifies Immune-Related Correlates of Resistance. Clin Cancer Res. 2013;19(10):2775−2786.

18. Patani N, Dunbier AK, Anderson H, Ghazoui Z, Ribas R, Anderson E, et al. Differences in the transcriptional response to fulvestrant and estrogen deprivation in ER-positive breast cancer. Clin Cancer Res. 2014;20(15):3962−3973.

19. Arthur LM, Turnbull AK, Webber VL, Larionov AA, Renshaw L, Kay C, et al. Molecular changes in lobular breast cancers in response to endocrine therapy. Cancer Res. 2014;74(19):5371−5376.

20. Turnbull AK, Kitchen RR, Larionov AA, Renshaw L, Dixon JM, Sims AH. Direct integration of intensity-level data from Affymetrix and Illumina microarrays improves statistical power for robust reanalysis. BMC Med Genomics. 2012;5:35.

21. Du P, Kibbe WA, Lin SM. lumi: a pipeline for processing Illumina microarray. Bioinformatics. 2008;24(13):1547−1548.

22. Yu G, He QY. ReactomePA: an R/Bioconductor package for reactome pathway analysis and visualization. Mol Biosyst. 2016;12(2):477−479.

23. Hong FX, Breitling R, McEntee CW, Wittner BS, Nemhauser JL, Chory J. RankProd: a bioconductor package for detecting differentially expressed genes in meta-analysis. Bioinformatics. 2006;22(22):2825−2827.

24. Likhite N, Warawdekar UM. A unique method for isolation and solubilization of proteins after extraction of RNA from tumor tissue using trizol. J Biomol Tech. 2011;22(1):37−44.

25. Coleman O, Henry M, Clynes M, Meleady P. Filter-Aided Sample Preparation (FASP) for Improved Proteome Analysis of Recombinant Chinese Hamster Ovary Cells. Methods Mol Biol. 2017;1603:187−194.

26. Rappsilber J, Ishihama Y, Mann M. Stop and go extraction tips for matrix-assisted laser desorption/ionization, nanoelectrospray, and LC/MS sample pretreatment in proteomics. Anal Chem. 2003;75(3):663−670.

27. Cox J, Mann M. MaxQuant enables high peptide identification rates, individualized p.p.b.-range mass accuracies and proteome-wide protein quantification. Nat Biotechnol. 2008;26(12):1367−1372.

28. Vizcaino JA, Csordas A, Del-Toro N, Dianes JA, Griss J, Lavidas I, et al. 2016 update of the PRIDE database and its related tools. Nucleic Acids Res. 2016;44(22):11033.

29. Haffner MC, Chaux A, Meeker AK, Esopi DM, Gerber J, Pellakuru LG, et al. Global 5-hydroxymethylcytosine content is significantly reduced in tissue stem/progenitor cell compartments and in human cancers. Oncotarget. 2011;2(8):627−637.

30. Barbie DA, Tamayo P, Boehm JS, Kim SY, Moody SE, Dunn IF, et al. Systematic RNA interference reveals that oncogenic KRAS-driven cancers require TBK1. Nature. 2009;462(7269):108−U122.

31. Lamb J, Crawford ED, Peck D, Modell JW, Blat IC, Wrobel MJ, et al. The connectivity map: Using gene-expression signatures to connect small molecules, genes, and disease. Science. 2006;313(5795):1929−1935.

32. Jang ER, Lim SJ, Lee ES, Jeong G, Kim TY, Bang YJ, et al. The histone deacetylase inhibitor trichostatin A sensitizes estrogen receptor alpha-negative breast cancer cells to tamoxifen. Oncogene. 2004;23(9):1724−1736.

33. Cao B, Li J, Zhu J, Shen M, Han K, Zhang Z, et al. The antiparasitic clioquinol induces apoptosis in leukemia and myeloma cells by inhibiting histone deacetylase activity. J Biol Chem. 2013;288(47):34181−34189.

34. Chen EY, Tan CM, Kou Y, Duan Q, Wang Z, Meirelles GV, et al. Enrichr: interactive and collaborative HTML5 gene list enrichment analysis tool. BMC Bioinformatics. 2013;14:128.

35. Ellis MJ, Suman VJ, Hoog J, Goncalves R, Sanati S, Creighton CJ, et al. Ki67 Proliferation Index as a Tool for Chemotherapy Decisions During and After Neoadjuvant Aromatase Inhibitor Treatment of Breast Cancer: Results From the American College of Surgeons Oncology Group Z1031 Trial (Alliance). J Clin Oncol. 2017;35(10):1061−1069.

36. Jankowitz RC, Oesterreich S, Lee AV, Davidson NE. New Strategies in Metastatic Hormone Receptor-Positive Breast Cancer: Searching for Biomarkers to Tailor Endocrine and Other Targeted Therapies. Clin Cancer Res. 2017;23(5):1126−1131.

37. Miller WR, Larionov A. Changes in expression of oestrogen regulated and proliferation genes with neoadjuvant treatment highlight heterogeneity of clinical resistance to the aromatase inhibitor, letrozole. Breast Cancer Res. 2010;12(4):R52.

38. Zhang XHF, Giuliano M, Trivedi MV, Schiff R, Osborne CK. Metastasis Dormancy in Estrogen Receptor-Positive Breast Cancer. Clin Cancer Res. 2013;19(23):6389−6397.

39. Bartosh TJ. Cancer cell cannibalism and the SASP: Ripples in the murky waters of tumor dormancy. Mol Cell Oncol. 2017;4(1):e1263715.

40. Barkan D, Green JE, Chambers AF. Extracellular matrix: a gatekeeper in the transition from dormancy to metastatic growth. Eur J Cancer. 2010;46(7):1181−1188.

41. Wang G, Platt-Higgins A, Carroll J, de Silva Rudland S, Winstanley J, Barraclough R, et al. Induction of metastasis by S100P in a rat mammary model and its association with poor survival of breast cancer patients. Cancer Res. 2006;66(2):1199−1207.

42. Yang RX, Stocker S, Schott S, Heil J, Marme F, Cuk K, et al. The association between breast cancer and S100P methylation in peripheral blood by multicenter case-control studies. Carcinogenesis. 2017;38(3):312−320.

43. Peng C, Chen H, Wallwiener M, Modugno C, Cuk K, Madhavan D, et al. Plasma S100P level as a novel prognostic marker of metastatic breast cancer. Breast Cancer Res Treat. 2016;157(2):329−338.

44. Dakhel S, Padilla L, Adan J, Masa M, Martinez JM, Roque L, et al. S100P antibody-mediated therapy as a new promising strategy for the treatment of pancreatic cancer. Oncogenesis. 2014;3:e92.

45. Basse C, Arock M. The increasing roles of epigenetics in breast cancer: Implications for pathogenicity, biomarkers, prevention and treatment. Int J Cancer. 2015;137(12):2785−2794.

46. Crea F, Nur Saidy NR, Collins CC, Wang Y. The epigenetic/noncoding origin of tumor dormancy. Trends Mol Med. 2015;21(4):206−211.

47. Fang F, Turcan S, Rimner A, Kaufman A, Giri D, Morris LG, et al. Breast cancer methylomes establish an epigenomic foundation for metastasis. Sci Transl Med. 2011;3(75):75ra25.

48. Tsang JS, Vencken S, Sharaf O, Leen E, Kay EW, McNamara DA, et al. Global DNA methylation is altered by neoadjuvant chemoradiotherapy in rectal cancer and may predict response to treatment - A pilot study. Ejso. 2014;40(11):1459−1466.

49. De Smet C, Loriot A. DNA hypomethylation in cancer Epigenetic scars of a neoplastic journey. Epigenetics. 2010;5(3).

50. Pathiraja TN, Xi Y, Lee AV, Santen R, Gannon F, Kaipparettu B, et al. Estrogen Deprivation Results in Altered DNA Methylation Profile in Breast Cancer Cells - Role in Endocrine Resistance? Cancer Res. 2009;69(24):808s−808s.

51. Harris ME, Bohni R, Schneiderman MH, Ramamurthy L, Schumperli D, Marzluff WF. Regulation of histone mRNA in the unperturbed cell cycle: evidence suggesting control at two posttranscriptional steps. Mol Cell Biol. 1991;11(5):2416−2424.

52. Braunstein M, Liao L, Lyttle N, Lobo N, Taylor KJ, Krzyzanowski PM, et al. Downregulation of histone H2A and H2B pathways is associated with anthracycline sensitivity in breast cancer. Breast Cancer Res. 2016;18(1):16.

53. Zhu Z, Edwards RJ, Boobis AR. Increased expression of histone proteins during estrogen-mediated cell proliferation. Environ Health Perspect. 2009;117(6):928−934.

54. Sarkar S, Abujamra AL, Loew JE, Forman LW, Perrine SP, Faller DV. Histone Deacetylase Inhibitors Reverse CpG Methylation by Regulating DNMT1 through ERK Signaling. Anticancer Res. 2011;31(9):2723−2732.

55. Lee JH, Choy ML, Ngo L, Foster SS, Marks PA. Histone deacetylase inhibitor induces DNA damage, which normal but not transformed cells can repair. Proc Natl Acad Sci U S A. 2010;107(33):14639−14644.

56. Bolden JE, Shi W, Jankowski K, Kan CY, Cluse L, Martin BP, et al. HDAC inhibitors induce tumor-cell-selective pro-apoptotic transcriptional responses. Cell Death Dis. 2013;4:e519.

57. Ceccacci E, Minucci S. Inhibition of histone deacetylases in cancer therapy: lessons from leukaemia. Br J Cancer. 2016;114(6):605−611.

58. Masri S, Lui K, Phung S, Ye J, Zhou D, Wang X, et al. Characterization of the weak estrogen receptor alpha agonistic activity of exemestane. Breast Cancer Res Treat. 2009;116(3):461−470.

59. Aguilar H, Sole X, Bonifaci N, Serra-Musach J, Islam A, Lopez-Bigas N, et al. Biological reprogramming in acquired resistance to endocrine therapy of breast cancer. Oncogene. 2010;29(45):6071−6083.

60. Milosevic J, Klinge J, Borg AL, Foukakis T, Bergh J, Tobin NP. Clinical instability of breast cancer markers is reflected in long-term in vitro estrogen deprivation studies. BMC Cancer. 2013;13.

61. Simigdala N, Gao Q, Pancholi S, Roberg-Larsen H, Zvelebil M, Ribas R, et al. Cholesterol biosynthesis pathway as a novel mechanism of resistance to estrogen deprivation in estrogen receptor-positive breast cancer. Breast Cancer Res. 2016;18.

62. Coser KR, Wittner BS, Rosenthal NF, Collins SC, Melas A, Smith SL, et al. Antiestrogen-resistant subclones of MCF-7 human breast cancer cells are derived from a common monoclonal drug-resistant progenitor. Proc Natl Acad Sci U S A. 2009;106(34):14536−14541.

63. Alves CL, Elias D, Lyng M, Bak M, Kirkegaard T, Lykkesfeldt AE, et al. High CDK6 Protects Cells from Fulvestrant-Mediated Apoptosis and is a Predictor of Resistance to Fulvestrant in Estrogen Receptor-Positive Metastatic Breast Cancer. Clin Cancer Res. 2016;22(22):5514−5526.

